# The effect of Shifts in Fish Community Structure on PCDD/F temporal variability in common guillemot (Baltic Sea)

**DOI:** 10.1101/2025.04.14.648406

**Authors:** Yosr Ammar, Jens Olsson, Elena Gorokhova, Martin Sköld, Suzanne Faxneld, Jonas Hentati-Sundberg, Anne L. Soerensen

## Abstract

Persistent Organic Pollutants, specifically polychlorinated dibenzodioxins (PCDDs) and dibenzofurans (PCDFs) (PCDD/Fs) threaten marine ecosystems through biomagnification and bioaccumulation in top predators. Following environmental legislation introduction in the 1970s, PCDD/F concentrations declined in the Baltic Sea biota. However, previous studies signaled that this decline plateaued from the 1990s onward in common guillemot (*Uria aalge*) eggs. This period coincides with shifts in Baltic environmental conditions and food-web structure, including guillemot prey species. Here, we hypothesize that temporal variability of PCDD/Fs in guillemot eggs was driven by changing environmental conditions and food-web structure affecting prey availability. We identified phases in the temporal dynamic of fish community structure, linked them to PCDD/Fs rate of change, and identified potential environmental and anthropogenic drivers of these phases. Three distinct phases emerged: cod and herring dominance (1976–1986), sprat dominance (1987–2001), and stickleback increase (2002–2021). PCDD/F concentrations declined sharply during the first phase (-6.4% y^1^), plateaued during the second (0.27% y^1^), and resumed a decline during the third (-3.8% y^1^). The transition to a sprat-dominated phase increased guillemots’ dietary exposure to PCDD/Fs, contributing to the second phase plateau. Conversely, stickleback rise as a potential key prey species during the third phase may have facilitated the post-2002 decline. Shifts in fish community structure were driven by changes in temperature, salinity, zooplankton size, and fishing pressure. We conclude that both bottom-up (environmental conditions) and top-down (fisheries food-web dynamics) effects have cascaded through the ecosystem, reshaping fish community structure and influencing PCDD/F concentrations in guillemot eggs.

## Introduction

Over the past century, socioeconomic growth has driven major environmental changes leading to three interconnected global crises: climate change, biodiversity loss, and pollution (UNEP, 2023). Chemical pollution has become a prominent concern causing severe environmental and human health issues with the potential to alter the Earth system processes at different scales (Alpizar et al., 2019; Bernhardt et al., 2017; Naidu et al., 2021; Persson et al., 2022; Richardson et al., 2023). Among the most significant groups of pollutants, persistent organic pollutants (POPs) have been recorded in most ocean sediments, water columns, and biota (Alpizar et al., 2019; Luarte et al., 2023; Puri et al., 2023; Willis et al., 2022; Xie et al., 2022). POPs bioaccumulation in marine food-webs causes physiological, behavioral, reproductive, and survival changes in wildlife (Bergman, 1999; Helander et al., 2002; Roos et al., 2012; Vagi et al., 2021; Willis et al., 2022).

Previous studies suggested that climate change can increase biotas’ exposure, bioaccumulation, and biomagnification potential of POPs with major consequences for marine top predators (Alava et al., 2020, 2018; De Wit et al., 2022). Climate change and other anthropogenic pressures impact on ecosystems are interlinked with complex ecological changes such as changes in food-web structure and species habitats (Pinsky et al., 2020), resulting in a vast diversity of processes that can directly or indirectly affect levels and trends of POPs (De Wit et al., 2022). For instance, northward range-shift of species adapting to climate change and shifts in the trophic position of others may impact POPs concentrations in recipient populations (Borgå et al., 2022). Additionally, altered foraging patterns of seabirds and marine mammals could impact their exposure to POPs (De Wit et al., 2022). For example, diet changes in East Greenland polar bears resulted in consistently slower temporal declines in adipose concentrations of legacy POPs (McKinney et al., 2013). There is, however, still limited information on how these complex ecological dynamics, shaped by climate change, anthropogenic pressures, food-web shifts, and foraging patterns, might affect the levels of POPs in marine top predatory birds.

In this study, we focus on the two POP groups, polychlorinated dibenzodioxins (PCDDs) and polychlorinated dibenzofurans (PCDFs), collectively referred to as PCDD/Fs, and changes in their concentrations over time in the marine top predatory bird, the common guillemot (*Uria aalge*). PCDD/Fs are groups of hydrophobic and lipophilic organic compounds that bioaccumulate to different degrees in the food-web (Isosaari et al., 2004). The release of PCDD/Fs is regulated under the 1979 Convention on Long-Range Transboundary Air Pollution and included in the original list of 12 POPs under the Stockholm Convention adopted in 2001 (http://www.pops.int). PCDD/Fs are also listed as priority substances in the EU Water Framework Directive (2013/39/EU) and are regulated in the EU Industrial Emissions Directive (2010/75/EU). PCDD/Fs are mostly by-products of various industrial, incineration, and combustion processes. In the Baltic Sea, the main sources for the two are deposition, diffusion/resuspension, and riverine input (McLachlan and Undeman, 2020). Their emissions peaked in the 1970s when legislation was introduced resulting in continuous declines in the concentrations in Baltic Sea biota (Gauss et al., 2020; Soerensen et al., 2024).

Previous studies, based on Swedish monitoring data, have reported an unexpected deceleration in the rate of decline of PCDD/F concentrations in guillemot eggs and Atlantic herring (*Clupea harengus*) from the late 1990s to the end of 2010s (Miller et al. 2014; Wiberg et al. 2013; Miller et al. 2013; Nyberg et al. 2012). In herring, these patterns have been attributed to slower growth and decreasing lipid content of the fish (Miller et al., 2013). However, the causes of the plateau in concentrations in guillemot eggs have not yet been fully understood as emissions have continued to decrease, suggesting that other factors are at play. Notably, dioxin air emissions in Sweden have decreased by 73% since 1990 due to improved technologies in several industries (Naturvårdsverket, 2024). The steepest decrease occurred between 1990 and 2000, interestingly coinciding with the stagnation period in PCDD/F concentrations in guillemot eggs, further supporting the hypothesis that additional factors may influence the observed trends.

The guillemot population in the Baltic Sea exhibits a high degree of site fidelity with two-thirds of the population residing on the island of Stora Karlsö (Hentati-Sundberg and Olsson, 2016). The bird primarily feeds on clupeids, especially sprat *Spratus spratus* (over 90% of their diet) and herring, but other fish species such as stickleback are also included in their diet (Enekvist, 2003; Lyngs and Durinck, 1998; Morkūnė et al., 2016). Guillemots, as high trophic level organisms with localized residence, consistent fat content, and low inter-annual variability in PCDD/F concentrations, are an ideal sentinel species in the Baltic Sea (Miller et al., 2014).

Besides the impact of POPs such as PCDD/Fs, the Baltic Sea is one of the marine ecosystems most affected by climate change, eutrophication, and overfishing (Blenckner et al., 2021; Elmgren et al., 2015). In particular, the Baltic Sea has registered one of the highest increases in water temperature over the past century and was suggested as an ideal time machine for understanding climate-induced changes in the global coastal ocean (Belkin, 2009; Reusch et al., 2018; Rutgersson et al., 2014; The BACC II Author Team, 2015). The human impact has resulted in multiple regime shifts in the Baltic Sea ecosystem over time (Möllmann et al. 2009; Tomczak et al. 2022; Dippner, Möller, and Hänninen 2012; Alheit et al. 2005; Eklöf et al. 2020; Österblom et al. 2006; Casini et al. 2009). Regime shifts are characterized by abrupt changes in the food-web structure and function that persist over time and encompass multiple trophic levels (Conversi et al., 2015). In the late 1980s in the Baltic Sea, for example, a shift in the dominance of the top predator cod to the zooplanktivorous clupeid sprat cascaded across multiple trophic levels (Alheit et al., 2005; Möllmann et al., 2009, 2008; Tomczak et al., 2022). Since the early 2000s, another mesopredatory fish species, the three-spined stickleback, has increased dramatically in the Baltic Sea, likely as a result of favorable environmental conditions in combination with declines in their key predators (Olin et al., 2022; Olsson et al., 2019). Given that these structural changes in the food-web involve some of the guillemot’s primary prey species, we investigate whether they could be contributing to the observed plateau in PCDD/F concentrations in their eggs.

In this study, we hypothesize that, in addition to changes in environmental loads of PCDD/Fs, the temporal variability of PCDD/Fs in guillemot eggs is driven by changes in environmental conditions (climate change and eutrophication) and food-web structure. We test this hypothesis by analyzing the long-term dynamics of PCDD/F concentrations in guillemot eggs in combination with environmental data and fish community structure in the Baltic Sea during the past four and half decades (1975-2021). Understanding marine ecosystem dynamics requires acknowledging their complexity, where nonlinear relationships between species and their environment as well as feedbacks often lead to unexpected changes in ecosystem structure and functions (Levin, 1998; Scheffer et al., 2001). Hence, we first identify major structural changes in the offshore Baltic Sea fish community, which comprise the common guillemot prey species. Second, we link these to the temporal changes in the PCDD/F concentrations (including their toxic equivalent) in common guillemot eggs and discuss drivers of temporal variability in the ecosystem.

## Material and methods

### 1. Data description

**PCDD/F congeners (lipid weight, lw)** in guillemot eggs from the Swedish National Monitoring Program for Contaminants in Marine Biota covering the years 1968-2022 were used. The eggs were collected from the island of Stora Karlsö in the Central Baltic Sea (Figure 1) and the sampling and analysis methodology described in (Ammar et al., 2024a) The time series contains a data gap from 1996 to 2001 for all congeners except 2,3,4,7,8-Pentachlorodibenzofuran. Seven congeners were determined for PCDDs and ten for PCDFs (Supplementary Table 1; Figure S1). Each year, ten eggs were analyzed either individually or as homogenates (pooled samples). The analyses of PCDD/Fs were carried out at the Department of Chemistry, Umeå University. The extraction method is described by Wiberg et al., (1998), the clean-up method by Danielsson et al., (2005), and the instrumental analysis (GC-HRMS) by Liljelind et al., (2003). The time series are presented in Figure S1 and the data is available in Ammar et al., (2024b).

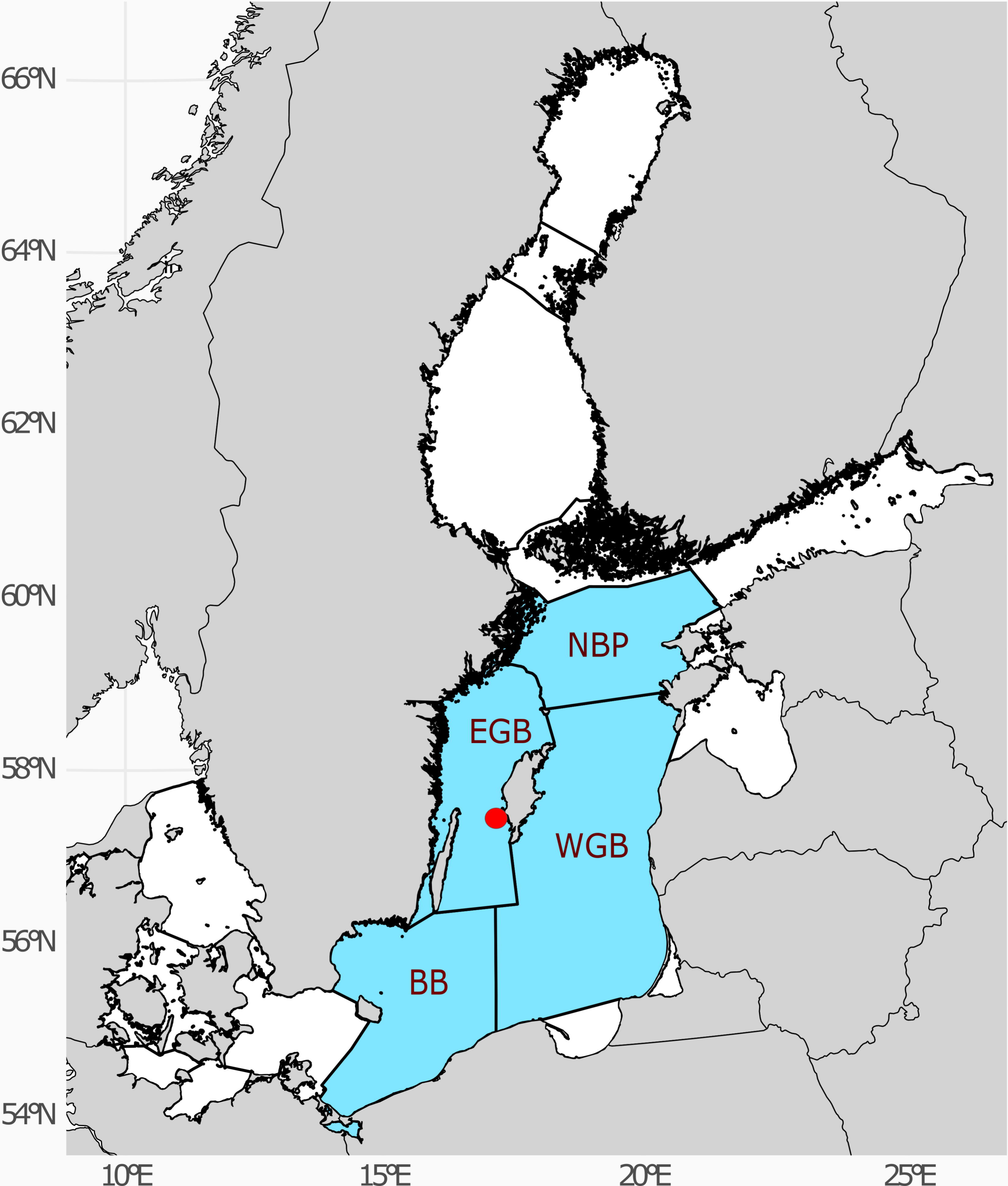

**Toxic Equivalent (TEQ) of PCDD/Fs or TEQ_PCDD/Fs_** (pg/g lw) were calculated using bird-specific Toxicity Equivalency Factors (TEFs) as recommended by Van Den Berg et al., (1998). These TEFs account for differences in sensitivity to PCDD/Fs between birds, mammals, and fish. By combining measured concentrations of congeners with their respective TEFs, the TEQ represents the total toxicity of a mixture. Between 1996 and 2001, TEQ_PCDD/Fs_ was calculated by quantifying 2,3,4,7,8-Pentachlorodibenzofuran and estimating the remaining congeners based on their typical ratios to 2,3,4,7,8-Pentachlorodibenzofuran, and applying the respective TEFs (Haglund et al., 2000).

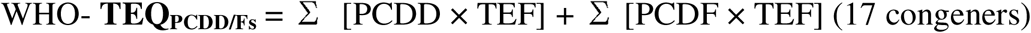

**Fish community** data describing the development and characteristics of the major offshore fish species in the Baltic Sea: Atlantic herring (*Clupea harengus*), European sprat (*Sprattus sprattus*), Atlantic cod (*Gadus morhua*), and three-spined stickleback (*Gasterosteus aculeatus*, Table 1) were used. As guillemot forages in the open offshore Baltic Sea (Hentati-Sundberg et al., 2018), we do not include benthic flatfish species in our analyses as there is no diet overlap between these and guillemots nor direct feeding (Morkūnė et al., 2016). We include data for the following stocks: Central Baltic herring, Baltic sprat, and Eastern Baltic cod (ICES, 2024) and stickleback of the offshore Baltic area (Bergström et al., 2015). Stock variables used for herring, sprat, and cod are recruitment (R: number of new individuals that survive to enter the adult population in a given time period) and spawning stock biomass (SSB: total weight of the mature portion of a fish population that is capable of reproducing). Fishing mortality (F: annual rate of the proportion of the fish population that is harvested or killed in fishing) is included as a pressure. These data were taken from the most recent report of the ICES Baltic Fisheries Assessment Working Group (ICES, 2024). For three-spined stickleback, only relative abundance estimates were available (RA: number of individuals per trawl hour) from the Baltic International Acoustic Survey sensu Bergström et al., (2015). The time series can be found in the appendix Figure S2.

**Table 1:** Detailed description of the data (predictor and response variables) used in the analysis time series covering the years 1976 to 2021. All data are averaged over all Central Baltic Sea Basins, except zooplankton, which excludes BB data (Figure 1). Data sources are: fish stocks from ICES Baltic Sea assessment (ICES, 2024), the Baltic International Acoustic Survey sensu (Bergström et al., 2015), zooplankton from ICES DOME database (https://dome.ices.dk/views/Zooplankton.aspx), and environmental data from SharkWeb database (https://sharkweb.smhi.se/).

| Group | Data | Full name |
| --- | --- | --- |
| Fish | Herring R | Central Baltic herring recruitment (Age 0) |
|  | Sprat R | Baltic sprat recruitment (Age 0) |
|  | Cod R | Eastern Baltic cod recruitment (Age 0) |
|  | Herring SSB | Central Baltic herring spawning stock biomass (tonnes) |
|  | Sprat SSB | Baltic sprat spawning stock biomass (tonnes) |
|  | Cod SSB | Eastern Baltic cod spawning stock biomass (tonnes) |
|  | Herring F | Central Baltic herring fishing mortality (tonnes) |
|  | Sprat F | Baltic sprat fishing mortality (tonnes) |
|  | Cod F | Eastern Baltic cod fishing mortality (tonnes) |
|  | Stickle RA | Stickleback relative abundance |
| Zooplankton | Zoo TZB | Summer (May-September) total zooplankton biomass (in mg/m<sup>3</sup>) |
|  | Zoo MS | Summer (May-September) zooplankton mean size (in µg/ind) |
| Environmental | WSAL | Winter surface (0-10m) salinity (PSU) |
|  | WBSAL | Winter bottom (>100 m in EGB, WGB, NBP, and >90 m in BB) salinity (PSU) |
|  | SuSST | Summer (June-August) sea surface (0-10 m) temperature (°C) |
|  | SpSST | Spring (March-May) sea surface (0-10 m) temperature (°C) |
|  | AuSST | Autumn (September-November) sea surface (0-10 m) temperature (°C) |
|  | SuSBT | Summer (June-August) sea bottom (>100m in EGB, WGB, NBP, and >90m in BB) temperature (°C) |
|  | DIP | Winter (December-February) Dissolved inorganic phosphorus (µmol/l) |
|  | DIN | Winter(December-February) Dissolved inorganic nitrogen (µmol/l) |
|  | Oxygen | Winter(December-February) deep-water (>100 m in EGB, WGB, NBP and >90 m in BB) dissolved oxygen (ml/l) |
|  | Secchi | Summer (June-August) Secchi depth (m) |

**Zooplankton** abundance data for May to September (growth season) from 1976 to 2021, which includes the spring and summer blooms comprising prey for the fish community, were retrieved from the ICES DOME database (https://dome.ices.dk/views/Zooplankton.aspx) as part of the Baltic Data Flows project (https://balticdataflows.helcom.fi). The dataset was curated to remove erroneous entries using an MSTS workflow script (Labuce and Gorokhova, 2023). This script was also applied to calculate (i) total zooplankton biomass (Zoo TZB; mg/m^3^) using individual wet weights (Hernroth, 1985); for species not included in this list, either measured or calculated individual weights based on length measurements were used, and (ii) mean size of a zooplankter in the community (Zoo MS; µg/ind.) based on the abundance and Zoo TZB values for each sampling occasion and calculating monthly averages that were further used to calculate summer means (Gorokhova et al., 2016). The data from the BB (Figure 1) were excluded due to a substantially shorter time series. The time series are presented in Figure S3.

**Environmental variables** were extracted from the SharkWeb database (https://sharkweb.smhi.se/; Table 1). We use the annual average of winter surface salinity (WSAL), winter bottom salinity (WBSAL), seasonal average of sea surface temperature: summer (SuSST), spring (SpSST), autumn (AuSST) as well as summer sea bottom temperature (SuSBT), the winter average dissolved inorganic nitrogen (DIN) and phosphorus (DIP), the average winter deep-water dissolved oxygen (Oxygen) and the summer average of Secchi depth (Secchi). The variables used in the analysis are commonly employed in the literature (e.g. Tomczak et al. 2022) and represent the major characteristics of the Baltic ecosystem and its changes related to climate change and eutrophication. The time series can be found in Figure S3.

### 2. Data processing

PCDD/F concentrations time series that contained more than ten years of data below the limit of quantification (LOQ) were excluded (Supplementary Table 1; Figure S1). Fish time series were provided from the data sources as individual time series for each variable. Missing values in the stickleback time series were imputed by interpolating between neighboring observations as previously used (maximum gap of 3 years). Each time series was then normalized (min-max scaling). Zooplankton time series were normalized by min-max scaling and then imputed by interpolating between neighboring observations in each basin (maximum gap of 3 years). These data were averaged for the Central Baltic Sea area (in this case NBP, WGB, and EGB, **Error! Reference source not found.**). Environmental time series from many sampling stations were averaged for each HELCOM subbasin (NBP, WGB, EGB, and BB). Missing environmental data were imputed by interpolating between neighboring observations in each basin (maximum gap of 3 years). The data were then averaged for the whole Central Baltic Sea area (**Error! Reference source not found.**). This was done to avoid over-representing conditions in one of the basins over others. Finally, each time series was normalized (min-max scaling).

### 3. Identification of phases in the fish community structure

To distinguish shifts in the general patterns of the fish community structure, we identified phases as periods with similar fish community compositions. We used the variables recruitment and spawning stock biomass (SSB) for sprat, cod, and herring, and relative abundance for stickleback (as outlined in Table 1) to characterize the fish community structure. To classify these phases, we applied a Gaussian mixture model using the *mclust* R package (Scrucca et al., 2016), which allows the number of clusters to be determined objectively via Bayesian Information Criterion (BIC). Unlike distance-based clustering, GMM assigns data points to clusters probabilistically, estimating the likelihood of each observation belonging to different distributions rather than relying on explicit similarity measures.

### 4. PCDD/Fs rate of change analysis relative to fish community phases

With the introduction of emission legislation in the 1970s, we anticipated a steady decrease in the PCDD/F concentrations in the Baltic Sea and its biota, assuming the environment remained otherwise stable. To identify deviations from this pattern, we estimate the rate of change of PCDD/F and TEQ_PCDD/Fs_ time series for each phase, by fitting linear splines with knots placed at the phase transitions identified by the GMM-derived in the fish community shifts. We do not include an average for all PCDD/Fs rate of change by phase as TEQ_PCDD/Fs_ takes into account all PCDD/Fs toxicity. We also calculate the overall rate of change for TEQ_PCDD/Fs_ time series over the whole time period studies by fitting linear splines to log-transformed series.

### 5. Analysis of predictors for each phase

To identify the drivers associated with each of the defined phases in fish community structure (drivers of differences between phases), we used Boosted Regression Trees (BRT). BRT is a non-parametric regression technique that combines a large number of simple decision trees by sequential fitting each new tree to the residuals of the previous ones without explicit assumptions about the functional form of the relationships between input variables (e.g., environmental variable) and the target variable (fish community structure, Elith et al., 2008; Jouffray et al., 2019). This method can fit nonlinear relationships, which allows to understand complex interactions in each phase and does not explicitly model temporal autocorrelation. This approach aligns with the understanding that BRTs can implicitly account for certain types of dependence (Elith et al., 2008). To run the BRT analysis and identify the relative influence of predictors on each phase, we used the R package *ggBRT* and followed the routine described by Jouffray et al., (2019).

Prior to the analysis, we used a multi-step approach to identify the candidate predictors of the fish community structure (responses) and to ensure variables free from excessive collinearity. We conducted a Variance Inflation Factor (VIF) analysis, setting a threshold of 5 to eliminate multicollinearity among the predictors. To further refine our selection, we performed a pairwise correlation analysis, removing variables with a correlation coefficient greater than 0.7 (Figure S4) following Jouffray et al. (2019).

To run the BRT analysis, we use the *gbm.step*() function with default parameters: a tree complexity of 5, a learning rate of 0.001, and a bag fraction of 0.75. These parameters allow consistency and comparability while balancing complexity and generalization between models. Alternative parameter values were tested and did not show significant differences. Cross-validation performance metrics included deviance, correlation, Area Under the Curve (AUC), and the percentage of explained deviance and highlighted robust models. For fish community structure, cross-validation explained deviance ranged from 72.43% to 80.16%, with high correlations (0.88–0.95) and AUC values (0.97–1), indicating robust predictive accuracy. All cross-validation results description is found in Supplementary Table 1.

All analyses were performed using R language and environment (R Core Team, 2022).

## Results

### 1. Phases in the fish community structure

Figure 2 shows the time series used to assess the fish community structure of the Baltic offshore food-web. Cod and herring spawning stock biomass (SSB) and recruitment (R) decreased during the late 1980s and early 1990s. In contrast, sprat spawning stock biomass (SSB) increased, peaking in the mid-1990s, while recruitment fluctuated over time. Data on the relative abundance (RA) of stickleback suggests that this species only occurred in very low population numbers until the early 2000s when a sharp increase occurred followed by a steady upward trend despite large interannual variability.

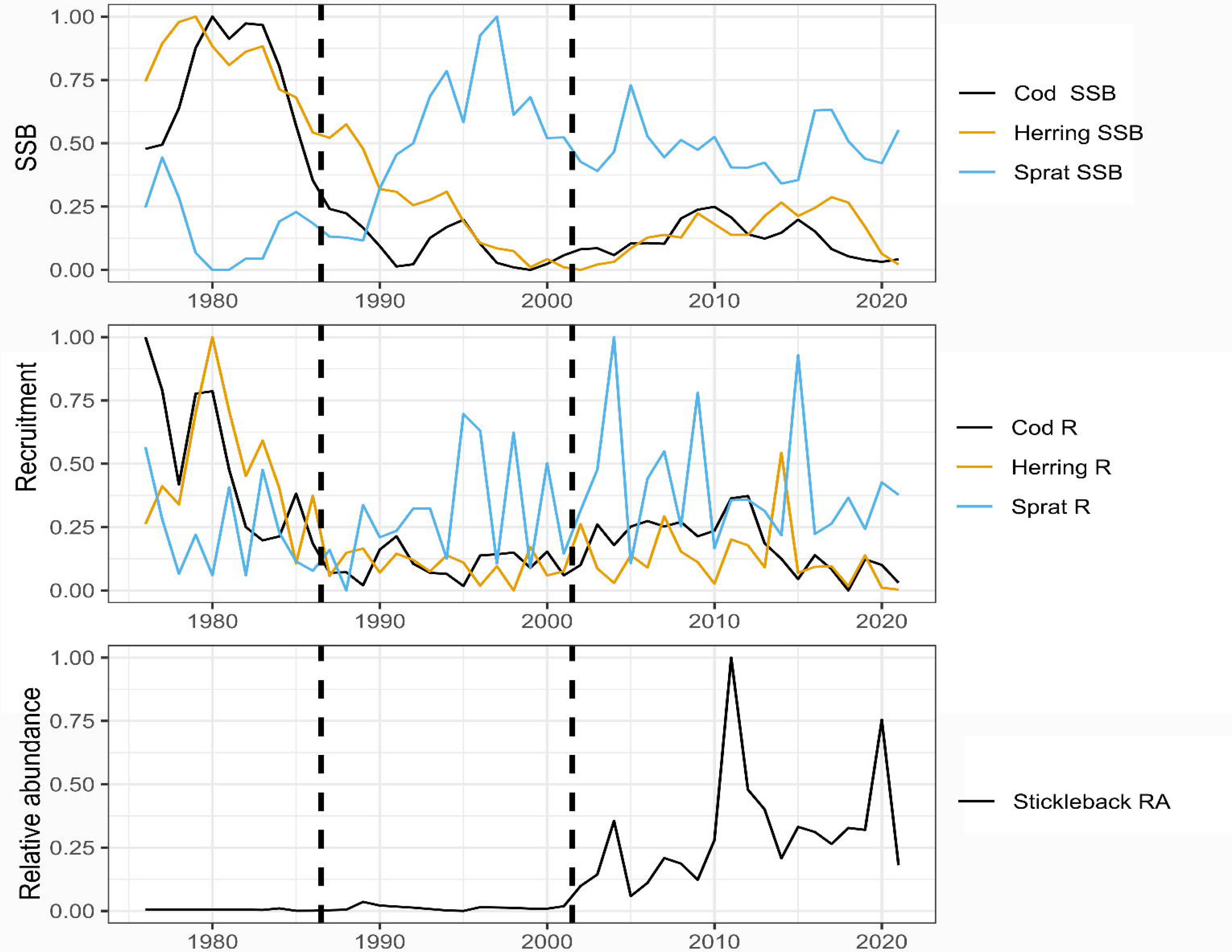

When analyzing changes in the fish community structure (GMM analysis), we identified three phases: 1976-1986, 1987-2001, and 2002-2021 (Figure 2, Figures S 4 & 5). Cod and herring recruitment, sprat spawning stock biomass (SSB), and relative stickleback abundance primarily drove differences between clusters. Cod SSB was the most influential variable in separating clusters, followed by sprat SSB and cod recruitment. Notably, stickleback relative abundance increased markedly in the most recent phase compared to earlier phases. Variance estimates suggest the highest variability in cod recruitment and sprat SSB, particularly in the second and third phases.

Hence, the first phase is characterized by high SSBs of cod and herring, a relatively low SSB of sprat and very low stickleback relative abundance. The second phase includes a sharp decrease in the SSBs of cod and herring and a rapid increase in the SSB of sprat. The last phase is characterized by a high SSB of sprat, low SSB of herring and cod, and a distinct increase in the relative abundance of stickleback.

### 2. Annual PCDD/F concentrations rate of change during three phases

The rate of change in concentrations of TEQ_PCDD/Fs_ for the entire time series is -2.8% yL¹. Using the identified phases of the fish community structure as change points (1987 and 2002); we calculated the rate of change in concentrations of TEQ_PCDD/Fs_ and individual PCDD/Fs in guillemot eggs (Figure 3). The annual rate of change in TEQ_PCDD/Fs_ was - 6.4% yL¹ during 1976–1986 (cod- and herring-dominated phase), followed by a near plateau with a rate of 0.27% yL¹ from 1987–2001 (sprat-dominated phase), and a resumed decline of -3.8% yL¹ between 2002–2021 (stickleback increase phase; Figures 3 & 4). Most individual congeners followed a similar pattern where the highest rate of decrease occurred in the first phase, followed by a lower decrease or plateau (no significant change) in the second phase. In the third phase, the rate of decrease for most congeners has picked up but is still slower than in the first phase. Exceptions to these patterns are seen for 2,3,7,8-Tetrachlorodibenzofuran, 1,2,3,7,8-Pentachlorodibenzofuran and 1,2,3,4,6,7,8,9-Octachlorodibenzo-p-dioxin. The individual time series and congener-specific differences are shown in Figure S1.

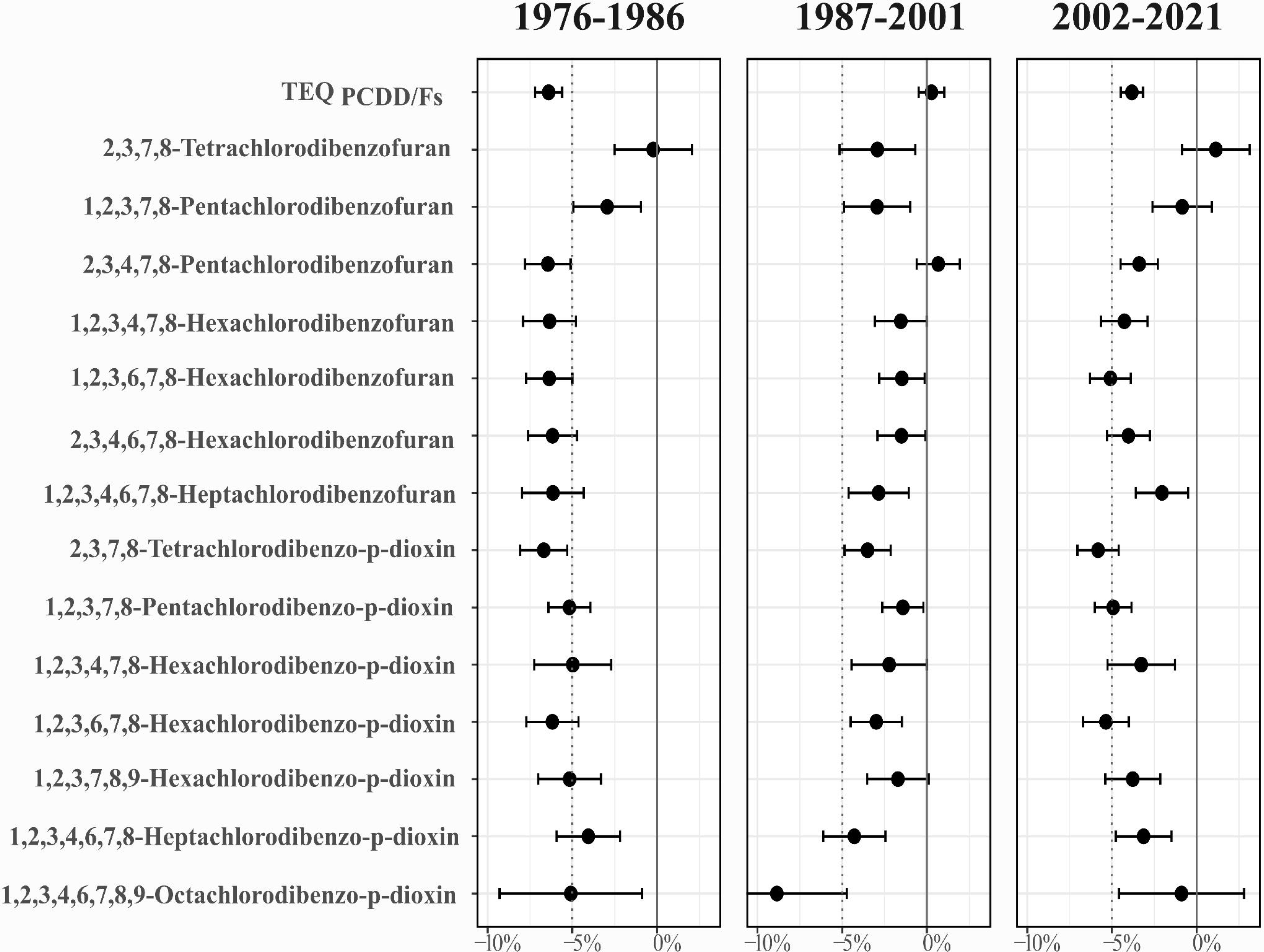

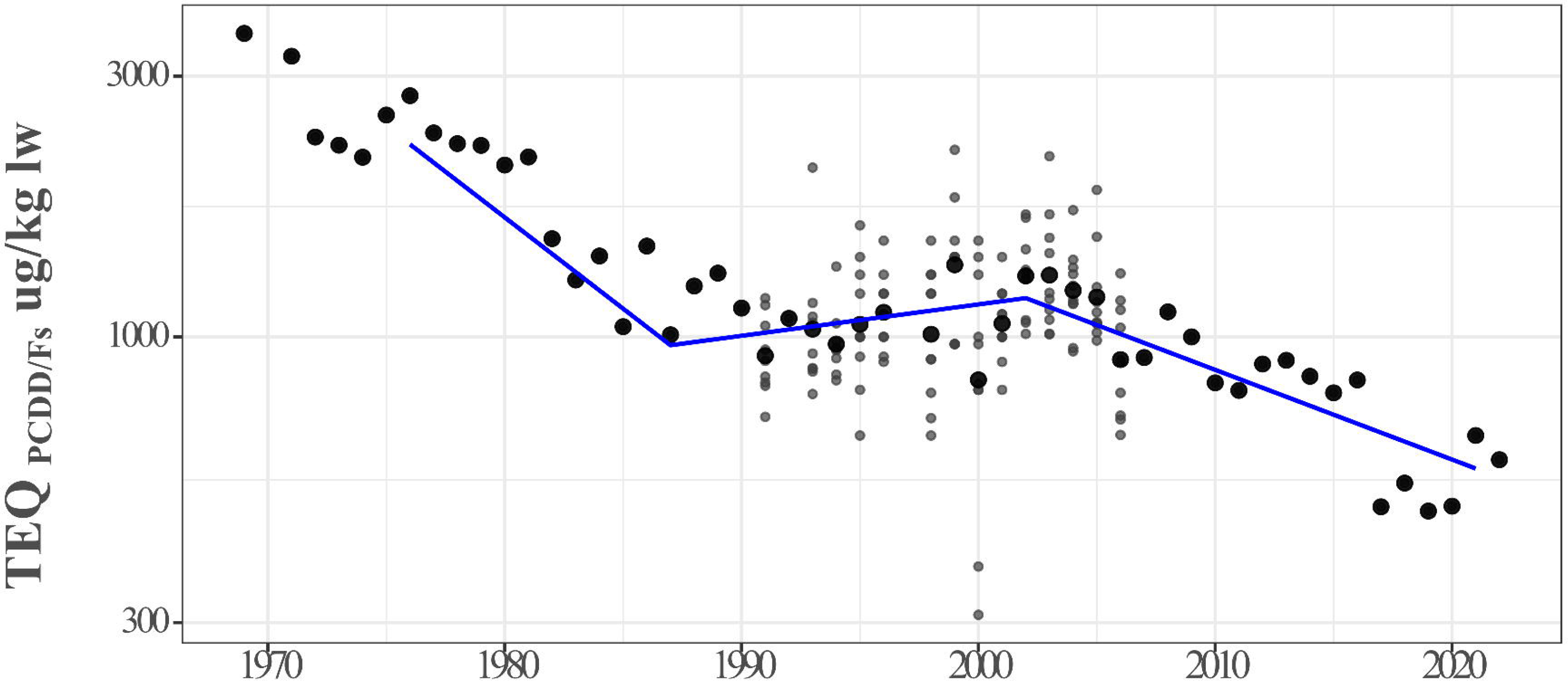

### 3. Predictors of change in the Baltic fish community structure

We identified the relative influence of potential predictors for each of the three fish community phases using BRT (Figure 5). We define a potential predictor as a variable with a relative influence on the specific phase above what could be expected by chance. During the first phase (1976-1986), the BRT suggested summer sea surface temperature (SuSST), herring fishing mortality (F), winter surface salinity (WSAL), and zooplankton mean size (Zoo MS) as potential predictors influencing the fish community structure. Temperature and herring fishing mortality exhibited negative relationships with the first phase suggesting that these variables were low when cod and herring dominated the fish community and PCDD/Fs were rapidly decreasing. On the contrary, winter salinity and zooplankton mean size showed positive relationships with the first phase in turn indicating that the fish community was associated with high salinity and high contribution of large-sized zooplankton. The predictors identified for the second phase (1987-2001) were herring and cod fishing mortality (F) (both with a negative relationship). This suggests that elevated fishing mortality on herring and cod are linked to the fish community structure during this phase, coinciding with a decline of cod and herring SSBs and an increase of sprat SSB, and a concurrent stagnation of PCDD/Fs. During the last phase (2002-2021), elevated summer sea surface temperature (SuSST, positive relationship) was identified as a potential predictor of the fish community structure, while zooplankton mean size (Zoo MS), dissolved inorganic nitrogen (DIN), oxygen, and cod fishing mortality (F) also had a relative influence above what is expected by chance and exhibiting negative relationships. This phase marked by higher stickleback and sprat abundance, lower cod and herring abundances, and an again increased decline in PCDD/Fs, hence appears to be influenced by higher water temperatures, reduced eutrophic conditions, smaller zooplankton, and lower cod fishing mortality than in the previous phases.

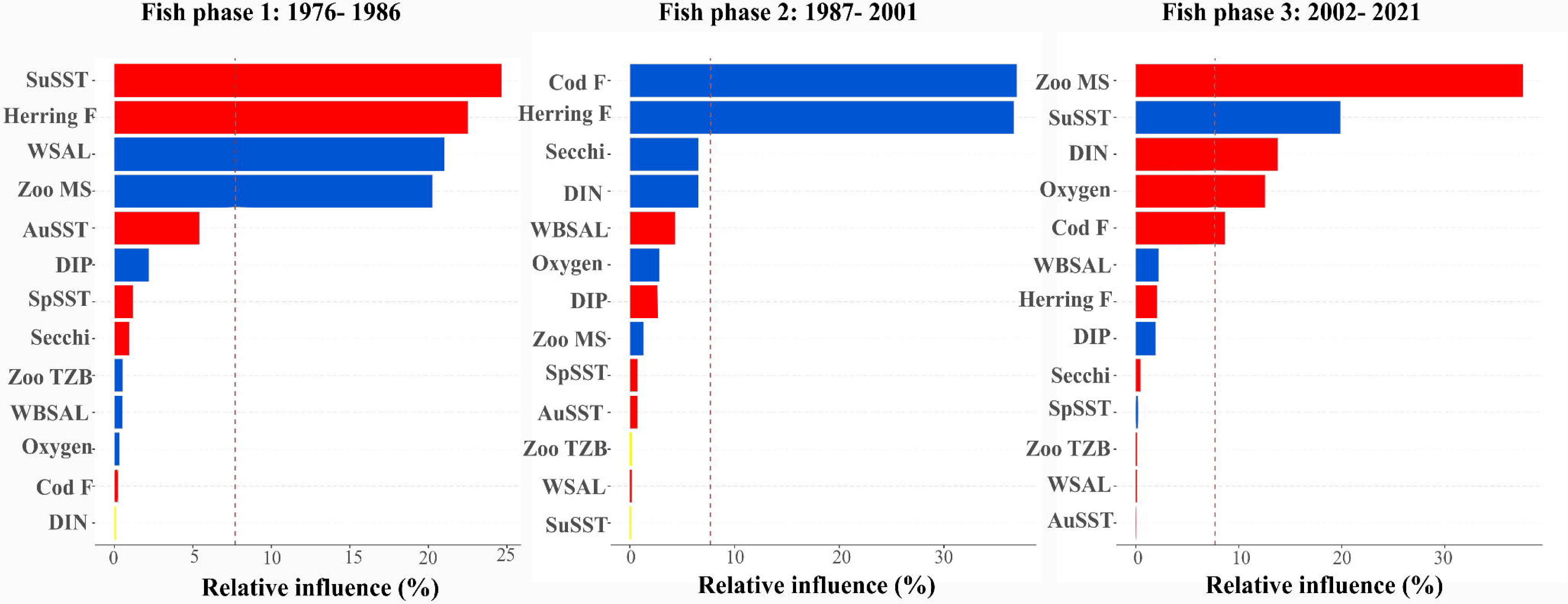

## Discussion

In this study, we investigate the environmental conditions and food-web structural changes associated with long-term PCDD/F trends in common guillemots in the Baltic Sea over the past four decades. Earlier observations indicated that while the initial decline in PCDD/Fs concentrations - driven by the emission control measures – was substantial, this trend plateaued during the 1990s, raising concerns about the Baltic Sea recovery and long-term environmental implications (Wiberg et al., 2009).

We hypothesized that changes in the offshore fish community structure affected PCDD/F concentrations in guillemot eggs, resulting in higher-than-expected levels. Our findings confirm that the initial decline of PCDD/F concentrations (-6.4% yL¹ during 1976-1986) plateaued during the 1990s (0.27% yL¹ during 1987-2001), consistent with previous reports (Miller et al. 2014; Wiberg et al. 2013; Nyberg et al. 2012). However, this stagnation was temporary, as the decline resumed at a moderated rate (-3.8% yL¹ during 2002-2021). Below, we explore the potential causes underlying these temporal changes and discuss their implications for future management and policy interventions.

### 1. Shifts in the offshore fish community structure of the Central Baltic Sea

To the best of our knowledge, this is the first study to analyze the offshore Central Baltic Sea fish community as a whole, including cod, herring, sprat, and three-spined stickleback, and to identify the most recent distinct phase (2002-2021) in its structure. Two shifts were found in the fish community dividing the study period into three distinct phases: 1976-1986, 1987-2001, and 2002-2021. The shift from the first to the second phase mirrors the well-documented late 1980s regime shift from a dominance of the top predatory fish species cod to the clupeid sprat in the Central Baltic Sea (Möllmann et al., 2009; Tomczak et al., 2022). Cod, which previously exerted a top-down control on sprat (Casini et al. 2009), suffered a dramatic biomass decline, resulting in a shift in the food-web dynamic from a benthic to a pelagic-dominated state (Tomczak et al., 2022). The most recent, third phase, beginning in 2002, is characterized by a sharp increase in the relative abundance of the stickleback alongside high sprat abundance and declines of cod and herring biomasses (Figure 2). Stickleback has been suggested to play an important role in the Baltic offshore pelagic food-web (Jakubavičiūtė et al., 2017; Olin et al., 2022), estimated to contribute to 10% of the pelagic fish biomass in the Baltic Proper and 40% of the estimated sprat biomass between 2011 and 2014 (Olsson et al., 2019).

### 2. Bottom-up impacts of fish community shifts on PCDD/F concentrations in guillemot eggs

By applying the three identified fish community phases to the guillemot time series, we observed phase-dependent changes in PCDD/F concentrations. We observed distinct variations in the rate of decrease across phases, with a steep decline in the first phase, a plateau during the second phase for TEQ_PCDD/Fs_ and most congeners, followed by an accelerated decline in the third phase. However, a few congeners deviated from this trend, particularly those known to exhibit distinct environmental behaviors compared to other PCDD/F congeners (Haglund et al., 2000). While this analysis does not determine the precise timing of shifts in PCDD/F concentration trends, it provides strong evidence of phase-dependent changes in concentration patterns, highlighting the role of fish community dynamics in modulating contaminant levels in guillemot eggs.

As suggested from our results, substantial changes in prey availability, driven by shifts in fish community composition, can cascade through the food-web, influencing PCDD/F concentrations in the guillemot population. Specifically, changes in the biomass of sprat, the main prey species of guillemot (Enekvist, 2003; Lyngs and Durinck, 1998; Morkūnė et al., 2016), might be an important driver for the observed plateau in decrease in PCDD/F concentrations in the eggs of the bird. The sprat stock, especially in terms of spawning stock biomass (SSB), increased dramatically in the 1990s (Figure 2). Previous research established that changes in the sprat population have had cascading effects on the common guillemots influencing chick body mass (Österblom et al., 2006) and fledging success (Kadin et al., 2012) but this is the first time this shift has been shown to also affect the accumulation of environmental contaminants.

The increase in the sprat population in the early 1990s (second phase) intensified competition within the clupeid community (sprat and herring), leading to density-dependent growth decline for both species (Casini et al. 2006). Sprat weight-at-age dropped by 40% during 1992–1998 and has since then remained low, with interannual fluctuations showing no clear trend (ICES, 2024). This decline in sprat and herring condition also resulted in a lower fat content of these prey species for guillemots compared to earlier years (ICES, 2024; Røjbek et al., 2014). Feeding rates of guillemot chicks on clupeids, mainly sprat hence almost doubled in 1998 and 2002 compared to 1975 (Enekvist, 2003; Österblom and Olsson, 2002) suggesting that guillemots compensated for the low-quality food by increasing their feeding rates (Kadin et al. 2016). This behavior may have contributed to the plateau of the rate of decline in PCDD/F concentrations in the guillemot eggs, due to increased consumption of low-quality sprat, a species that is known to accumulate PCDD/Fs (Vuorinen et al., 2012, 2002). A similar pattern was observed in Baltic salmon, where slow growth, high food competition, and reduced fat content of prey species (sprat and herring) enhanced the biomagnification of POPs, including PCDD/Fs (Vuorinen et al., 2012, 2002). Hence, our results support the hypothesis proposed by De Wit et al., (2022), that altered foraging patterns in seabirds, such as the common guillemot, can influence their exposure to POPs, here PCDD/Fs.

During the third phase (2002-2021), sprat SSB continued to be high although it exhibited a slight decrease compared to the 1990s and the condition (weight-at-age) of the fish remained low (ICES, 2024). Concurrently, the stickleback population started to increase (Figure 5). There are indications that stickleback has become an important prey for adult guillemots during the past two decades (Carlsen 2022; Kadin et al. 2016, JHS pers. obs.). Although sticklebacks are generally smaller than sprat, which might make them less energetically efficient for guillemots to catch and consume, they are often found in high abundance near the surface making them easily accessible (Carlsen, 2022). Due to their shorter lifespan (three to four years, Bergström et al., 2015) compared to sprat (five to six years, Peck et al., 2012), stickleback may accumulate less PCDD/Fs during their lifetime than sprat. Additionally, if sticklebacks have higher fat content than sprat, guillemots may consume less fish, leading to a lower accumulation of PCDD/Fs, although no data is available to support this hypothesis. Consequently, a shift during the third phase towards a diet including more stickleback might reduce the exposure to PCDD/Fs for the guillemots. This requires further investigation, to compare PCDD/F concentrations in stickleback and sprat, matched with detailed information on the guillemot diet.

### 3. Environmental and anthropogenic pressures impact the Baltic ecosystem

High stocks of cod and herring characterized the first phase of the fish community structure. Key environmental predictors associated with this phase included low summer sea surface temperature, high salinity, larger mean size zooplankton, and low herring fishing (Figure 5). The cooler temperatures and higher salinity during this period likely created favorable conditions for larger copepods, such as *Pseudocalanus acuspes*, a primary food source for cod larvae and the preferred prey for herring (Möllmann et al., 2008, 2005). However, throughout the study period, sea surface temperatures steadily increased, while salinity decreased after the second phase as direct consequences of climate change (HELCOM, 2023). These shifts favored sprat and smaller copepods (e.g., *Acartia* sp.) over cod, the main predator of sprat (Alheit et al. 2005; Ojaveer and Kalejs 2010; Möllmann et al. 2008). Since the 1990s, zooplankton mean size has declined severely driven by predation pressure, shifts in phytoplankton composition, and decreasing salinity – all of which impacted the availability and quality of fish prey (Gorokhova et al., 2016; HELCOM, 2023). This reduction in zooplankton size reflects an increased contribution of small cladocerans (e.g., *Bosmina*) and rotifers (e.g., *Keratella*) at the expense of copepods, resulting in a decline in the mass-specific lipid content of zooplankton (Gorokhova, 2019). This reduction in prey quality likely constrained fish growth, as herring and sprat increasingly compete for food and space (Diekmann et al., 2010; MacKenzie et al., 2007).

In the second sprat-dominated phase, the increase in fishing mortality of both cod and herring contributed to the observed increase in sprat. High herring fishing pressure is a potential reason for the decrease in herring stock, alongside reduced growth and lower recruitment success of the species (Diekmann et al., 2010). Additionally, the cod fishery has been one of the main anthropogenic activities that have impacted the biomass of the eastern Baltic cod stock in turn having effects on the whole food-web of the central Baltic Sea (Tomczak et al. 2022; Casini et al. 2008; Möllmann et al. 2009).

The third and most recent phase of fish community structure in the Baltic Sea is characterized by a substantial increase in the stickleback population, alongside a relatively stable sprat stock. This phase is associated with smaller zooplankton size, lower dissolved inorganic nitrogen (DIN), and oxygen concentrations, reduced cod fishing mortality, and rising summer sea surface temperature. Stickleback thrived in the presence of high abundance of small-sized low-lipid zooplankton during this phase, particularly cladocerans, which became more dominant as a result of salinity and temperature changes (Lankov et al., 2010). Sticklebacks consume cladocerans at higher rates than herring and sprat, played a key role in reducing competition for food (Lankov et al., 2010), further contributing to their competitive advantage (Jakubavičiūtė et al., 2017; Olin et al., 2022). Warmer water temperatures have likely benefited their ingestion and growth rates, as they migrate from the coast to the open sea for feeding at the end of summer (Bergström et al., 2015; Kotterba et al., 2014; Lefébure et al., 2014, 2011). Although stickleback can tolerate eutrophic conditions, their growth improves in less eutrophic waters (Olin et al., 2022), and the recent decrease in DIN may have further facilitated their expansion. Finally, the collapse of cod stock, driven by both fishing pressure and decreasing salinity and oxygen levels from reduced inflows from the North Sea, further contributed to changes in fish community structure (Fonselius and Valderrama, 2003; Hansson and Viktorsson, 2023; Tomczak et al., 2022, Figure S6) and led to the closure of the cod fishery (ICES, 2024). Consequently, the decline of cod and herring stocks—both predators of stickleback—combined with temperature and salinity changes, created favorable conditions for the stickleback population to thrive (Donadi et al., 2024; Olin et al., 2022). This may have contributed to the continued restructuring of the food-web, further altering dietary exposure to contaminants.

### 4. Concluding remarks and management implications

Changes in food-web structure driven by climate change and anthropogenic pressures can cascade through multiple trophic levels, ultimately affecting bioaccumulation patterns in top predators, such as fish-eating birds. Our results suggest the importance of fisheries management approaches that account for both direct and indirect food-web effects as cascading interactions can impact ecosystem dynamics. When, for example, setting single species quotas for fish stocks (Österblom et al., 2006), it is necessary to consider their broader ecological consequences, including unintended impacts on contaminant transfer within the food-web. As demonstrated in our study, the abundance, growth, and condition of targeted fish stocks can counteract management efforts to reduce contaminant loads in the ecosystem, stressing the need for a more holistic ecosystem-based approach to fisheries and environmental management.

Unlike for humans and fish, environmental quality standards for PCDD/Fs in birds remain poorly defined, and specific threshold values for adverse effects in these species are still uncertain. Hence, we cannot directly assess the toxicological significance of the TEQ_PCDD/Fs_ values in guillemot eggs reported in this study (Figure 4). However, the presence of POPs in general can weaken food-web resilience, even at concentrations below established toxicity thresholds and in the absence of observable population declines (Baudrot et al., 2018; Wolff et al., 2019). It is therefore imperative to continue efforts to reduce PCDD/Fs contamination and adopt ecosystem-based approaches, to avoid the reversal of reduction measures and safeguard ecosystem resilience in the face of ongoing environmental changes.

Over the past 20 years, PCDD/Fs emissions have remained stable and low in Europe but have increased in Africa and Asia (Lei et al., 2021). Today, PCDD/Fs are found in 60% of Arctic and Antarctic marine sediments, and the concentrations of POPs in polar regions are comparable to those reported in the southern Baltic Sea (Kobusińska et al., 2020). Climate change can exacerbate exposure to POPs for biota in the environment through secondary sources of reemission, remobilization from sea ice, and increased run-off (Borgå et al., 2022). Furthermore, changes in food-web structure due to climate change and anthropogenic pressures can lead to prolonged exposure for biota, as demonstrated in this study. These trends raise questions about the future persistence of PCDD/Fs and POPs globally, their impacts on marine ecosystems and top predators, and the effectiveness of current mitigation efforts. Despite this, chemical pollution remains largely overlooked in global change analysis (Bernhardt et al., 2017). There is an urgent need for interdisciplinary approaches that integrate pollution with other major global crises (climate change and biodiversity loss) to better understand their cumulative impacts on ecosystems across different scales and develop efficient mitigation strategies.

## Supporting information

Supplementary Tables and FiguresS

## Acknowledgements

We acknowledge financial support from the Formas - a Swedish Research Council for Sustainable Development (Grant No: 2021-00942). We thank current and former colleagues at the Swedish Museum of Natural History and sample analysis laboratories for their contribution to the Swedish National Monitoring Programme for Contaminants in Marine Biota. The Swedish Environmental Protection Agency funds the Swedish National Monitoring Programme for Contaminants in Marine Biota.

## Data statement

All data used in this manuscript are open-access and can be accessed through the links provided in the manuscript.

