## Supplementary Tables and FiguresS for "The effect of Shifts in Fish Community Structure on PCDD/F temporal variability in common guillemot (Baltic Sea)"

### 1. Data: Time series description

*Supplementary Table 1: PCDD/Fs analyzed in guillemot eggs from Stora Karlsö station in the Baltic Sea. Data sources are: PCDD/Fs from the Swedish national monitoring program for marine biota (Ammar et al., 2024a). * indicates congeners excluded from the analyses.*

| **PCDD/Fs** (pg/g lw) | **Full name** |
|---|---|
| TCDF | 2,3,7,8-Tetrachlorodibenzofuran |
| TCDD | 2,3,7,8-Tetrachlorodibenzo-p-dioxin |
| PECDF1 | 1,2,3,7,8-Pentachlorodibenzofuran |
| PECDF2 | 2,3,4,7,8-Pentachlorodibenzofuran |
| PECDD | 1,2,3,7,8-Pentachlorodibenzo-p-dioxin |
| HXCDF1 | 1,2,3,4,7,8-Hexachlorodibenzofuran |
| HXCDF2 | 1,2,3,6,7,8-Hexachlorodibenzofuran |
| HXCDF3 | 2,3,4,6,7,8-Hexachlorodibenzofuran |
| HXCDF4* | 1,2,3,7,8,9-Hexachlorodibenzofuran |
| HXCDD1 | 1,2,3,4,7,8-Hexachlorodibenzo-p-dioxin |
| HXCDD2 | 1,2,3,6,7,8-Hexachlorodibenzo-p-dioxin |
| HXCDD3 | 1,2,3,7,8,9-Hexachlorodibenzo-p-dioxin |
| HPCDF1 | 1,2,3,4,6,7,8-Heptachlorodibenzofuran |
| HPCDF2* | 1,2,3,4,7,8,9-Heptachlorodibenzofuran |
| HPCDD | 1,2,3,4,6,7,8-Heptachlorodibenzo-p-dioxin |
| OCDF* | 1,2,3,4,6,7,8,9-Octachlorodibenzofuran |
| OCDD | 1,2,3,4,6,7,8,9-Octachlorodibenzo-p-dioxin |

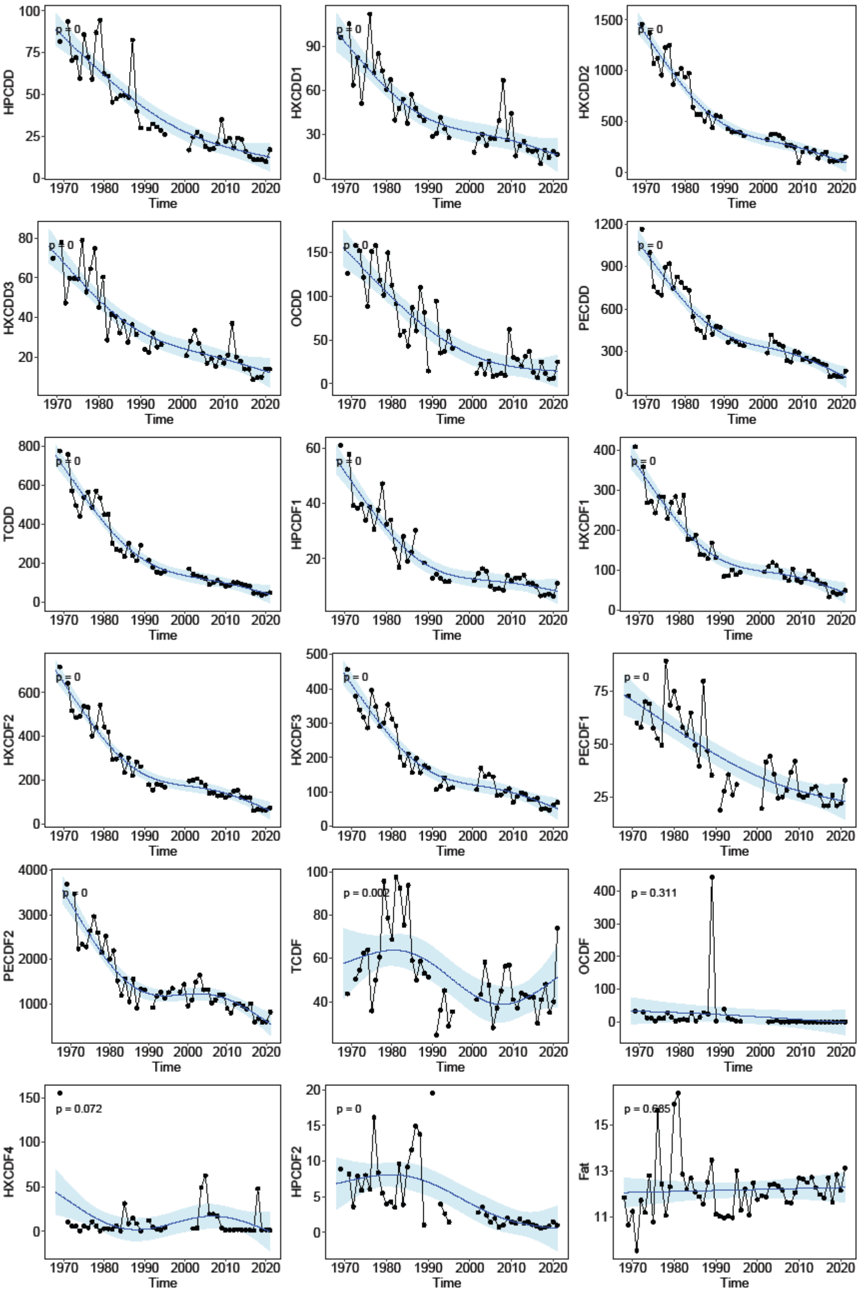


*Figure S1: Time series of PCDD/F concentrations in guillemot eggs between 1968 and 2021 as well as the fat % of the eggs with trend lines fitted using smoothing splines in a Generalized Additive Model (GAM). TCDF: 2,3,7,8-Tetrachlorodibenzofuran, TCDD: 2,3,7,8-Tetrachlorodibenzo-p-dioxin, PECDF1: 1,2,3,7,8-Pentachlorodibenzofuran, PECDF2: 2,3,4,7,8-Pentachlorodibenzofuran, PECDD: 1,2,3,7,8-Pentachlorodibenzo-p-dioxin, HXCDF1: 1,2,3,4,7,8-Hexachlorodibenzofuran, HXCDF2: 1,2,3,6,7,8-Hexachlorodibenzofuran, HXCDF3: 2,3,4,6,7,8-Hexachlorodibenzofuran, HXCDF4*: 1,2,3,7,8,9-Hexachlorodibenzofuran, HXCDD1: 1,2,3,4,7,8-Hexachlorodibenzo-p-dioxin, HXCDD2: 1,2,3,6,7,8-Hexachlorodibenzo-p-dioxin, HXCDD3: 1,2,3,7,8,9-Hexachlorodibenzo-p-dioxin, HPCDF1: 1,2,3,4,6,7,8-Heptachlorodibenzofuran, HPCDF2*: 1,2,3,4,7,8,9-Heptachlorodibenzofuran, HPCDD: 1,2,3,4,6,7,8-Heptachlorodibenzo-p-dioxin, OCDF*: 1,2,3,4,6,7,8,9-Octachlorodibenzofuran, OCDD: 1,2,3,4,6,7,8,9-Octachlorodibenzo-p-dioxin. * indicates congeners excluded from the analyses. Data sources are: PCDD/Fs from the Swedish national monitoring program for marine biota (Ammar et al., 2024a).*

Most congener time series in Figure S1 showed an overall downward trend with a plateau in the 1990s and 2000s. OCDD and TCDF are known to behave differently from other PCDD/F congeners (Haglund et al., 2000). HXCDF4, OCDF, and HPCDF2 time series were excluded from the analyses as values before 1995 are in majority below the limit of quantification (LOQ). The fat percentage (Fat %) fluctuated greatly before the mid-1980s, but became more stable with lower variability afterward.

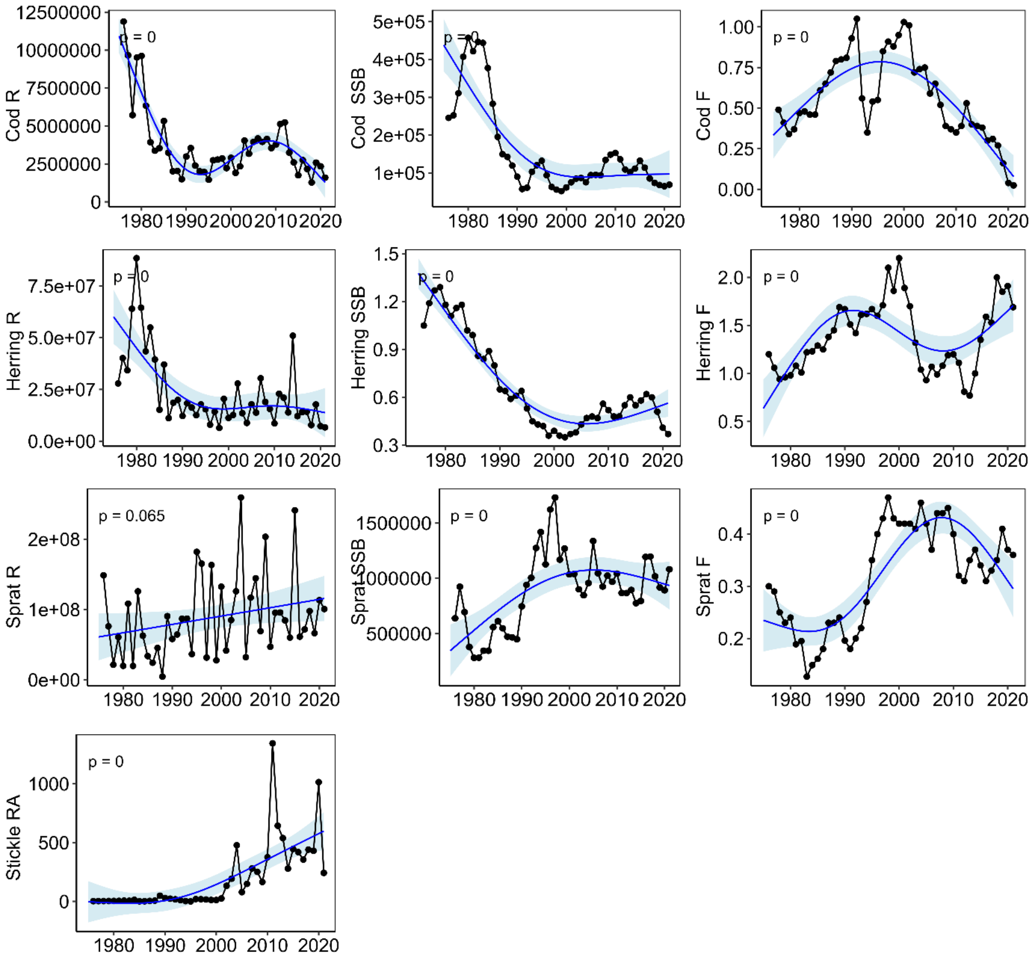


*Figure S2: Fisheries data time series with trend lines fitted using smoothing splines in a Generalized Additive Model (GAM) for cod, herring and sprat recruitment (R), spawning stock biomass (SSB) and fishing mortality (F) as well as stickleback relative abundance (Stickle RA).*

Cod recruitment decreased to a low value in the early 1990s, rose slightly in the late 2000s, and then decreased again until the end of the time series (Figure S2). Cod SSB peaked in the early 1980s dropped in the late 1980s and remained low thereafter. Cod fishing mortality increased gradually until the mid-1990s, dropped for a few years and in the 1980s, rose again to a high value in the early 2000s, then decreased gradually thereafter.

Herring recruitment peaked in the early 1980s, decreased on the mid-1980s, and remained at similar levels until the end of the time series (Figure S2). Herring spawning stock biomass decreased gradually until the early 2000s when it slightly increased, then it dropped again in the late 2010s. Herring fishing mortality increased gradually until the early 2000s, declined between the early 2000s and early 2010s, then increased again in the late 2010s.

Sprat recruitment was highly variable with an overall upward trend (Figure S2). Sprat spawning biomass increased gradually to reach the highest values in the 1990s, then decreased slightly in the 2000s, and

maintained a stable level afterwards. Sprat fishing mortality was low at the beginning of the time series, increased in the mid-1990s and remained high afterwards.
Stickleback relative abundance was null in the beginning of the time series and started increasing in the mid-2000s with high fluctuations (Figure S2).

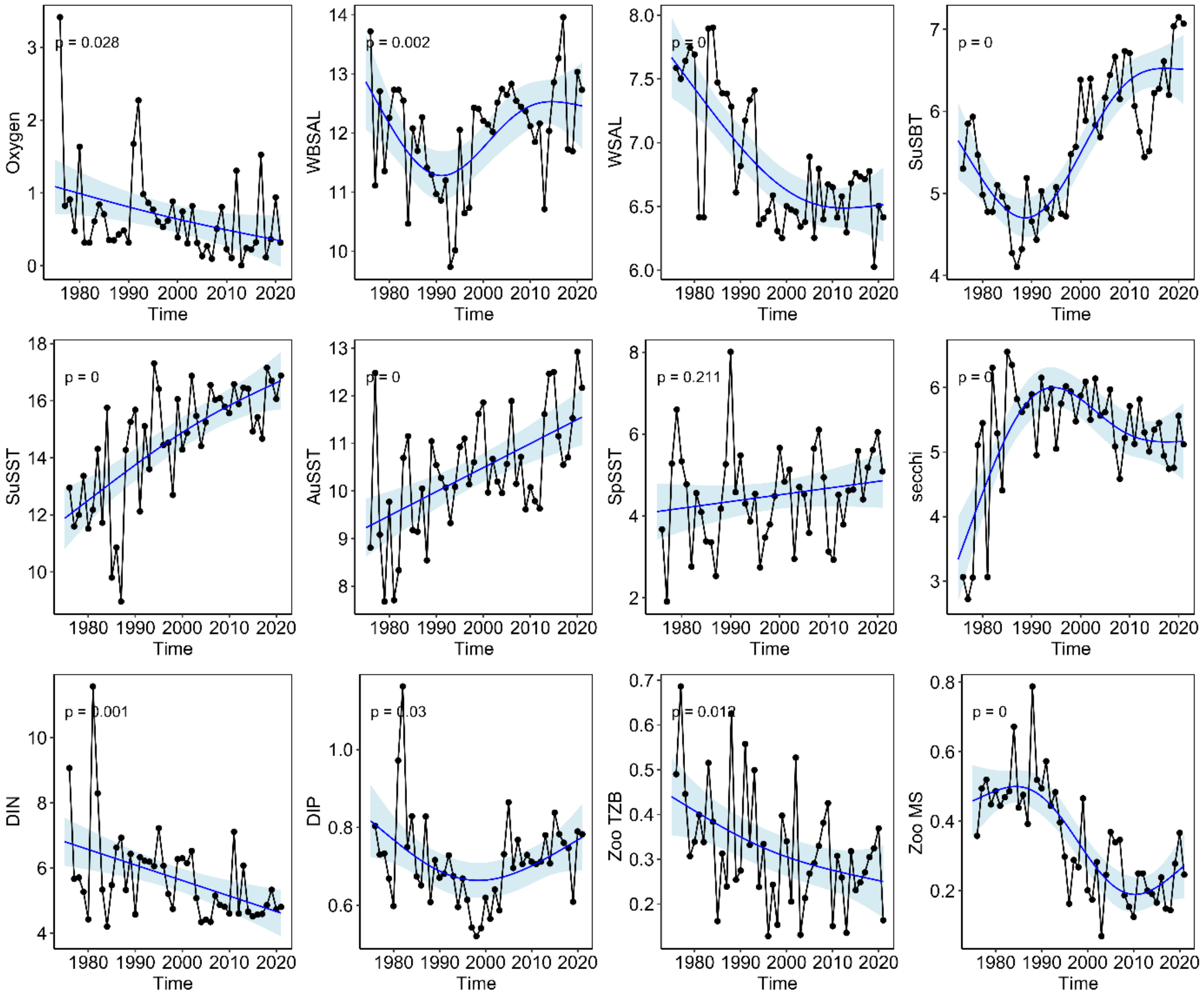


*Figure S3: Time series of environmental data with trend lines fitted using smoothing splines in a Generalized Additive Model (GAM): oxygen, winter bottom salinity (WBSAL), winter salinity (WSAL), summer bottom temperature (SuSBT), summer sea surface temperature (SuSST), autumn sea surface temperature (AuSST), spring sea surface temperature (SpSST), secchi depth, dissolved organic nitrogen (DIN), and dissolved organic phosphorus (DIP), and of zooplankton data: total zooplankton biomass (Zoo TZB) and zooplankton mean size (Zoo MS).*

Oxygen showed an overall downward trend with some fluctuations and close to hypoxic conditions towards the end of the time series (Figure S4). Winter bottom salinity declined from 13.5 to less than 10 psu, reached its lowest in the early 1990s, and increased in the early 2000s around 12.5 psu. Winter salinity fluctuates with a downward trend from around 7.5 to 6.5 psu. Summer bottom temperature decreased in the beginning of the time series with the lowest values found in the late 1980s, then increased gradually until the end of the time series. Summer, autumn and spring sea surface temperature had an overall upward trend with high fluctuations with respective 6 °C, 3 °C and 1 °C increase between the start and end of the time series. Secchi depth varies between 3 and 6 meter until the early 1990s then slowly decreases and fluctuates between 5 and 6 meters throughout the rest of the time serie. Dissolved inorganic nitrogen had an overall downward trend. Dissolved inorganic phosphorus showed a downward trend until the early 2000s then started increasing slightly.

Total zooplankton biomass showed an overall downward trend with high fluctuations. Zooplankton mean size decrease starting from the early 1990s.

### 2. Phases identification

GMM analysis identified three distinct phases in the fish community structure using a **VVI model** (diagonal covariance with varying volume and shape, Figure S4), as selected based on Bayesian Information Criterion (BIC). The optimal model included **three clusters** with respective sample sizes of **11, 15, and 20 observations** (Figure S5). The log-likelihood of the final model was **-3949.418**, with a **BIC of -8090.268** and an Integrated Complete Likelihood (ICL) of **-8090.337**, indicating a stable classification. The centroids of the clusters revealed clear temporal shifts, with the mean years for each phase centered around **1981, 1994, and 2011**, respectively. Classification uncertainty was minimal, as reflected in the cluster membership probabilities, where each observation was strongly assigned to one cluster (posterior probabilities close to 1).

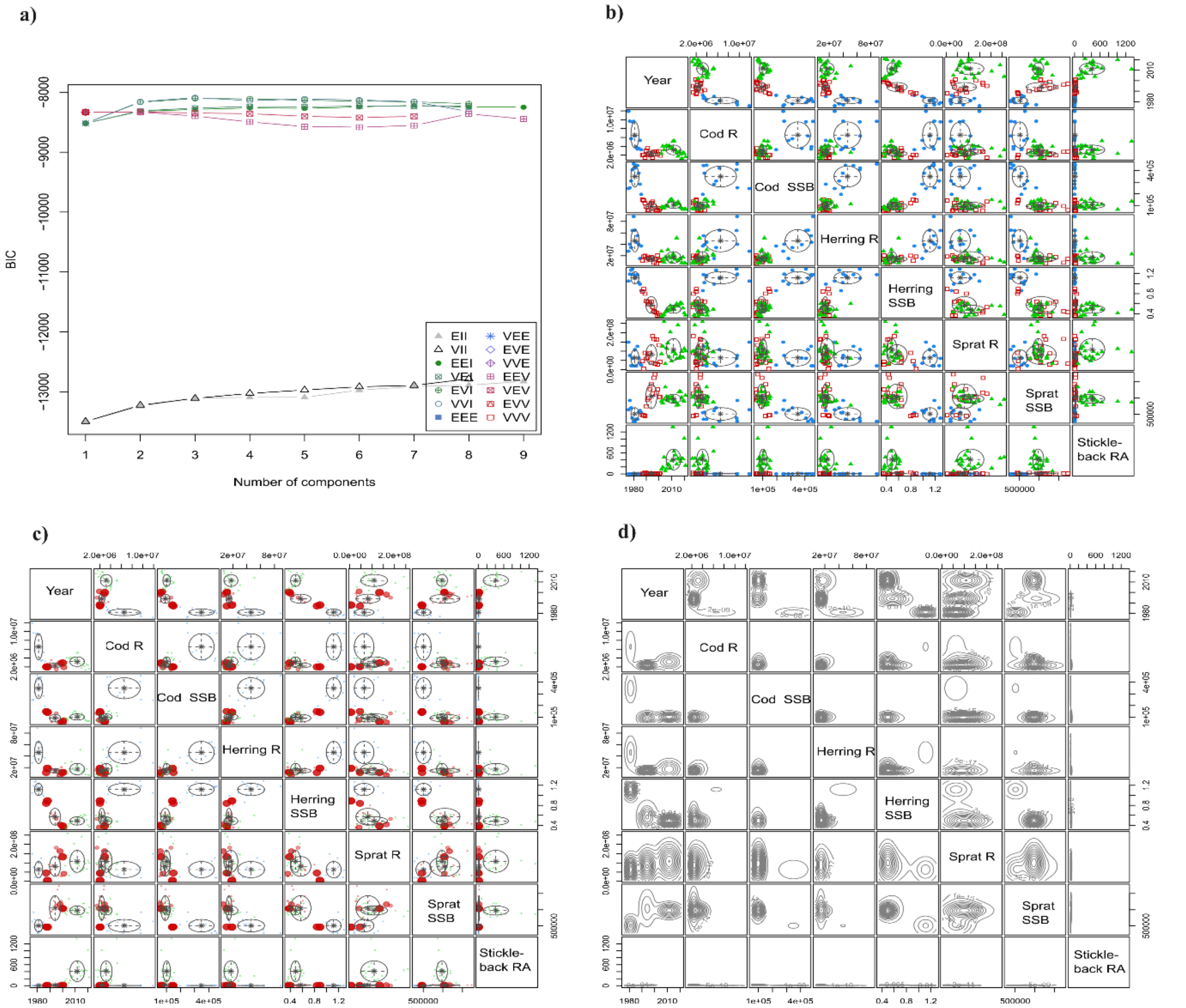


*Figure S 4: Model-based clustering analysis: a) Bayesian Information Criterion (BIC) showing model fit for different numbers of clusters, used for model selection with the chosen VVI model; b) Classification of data points into clusters based on the selected model; c) Uncertainty of classification, displaying the posterior probability of each data point belonging to a particular cluster; d) Density of the data points within each cluster, illustrating the distribution of data across different groups.*

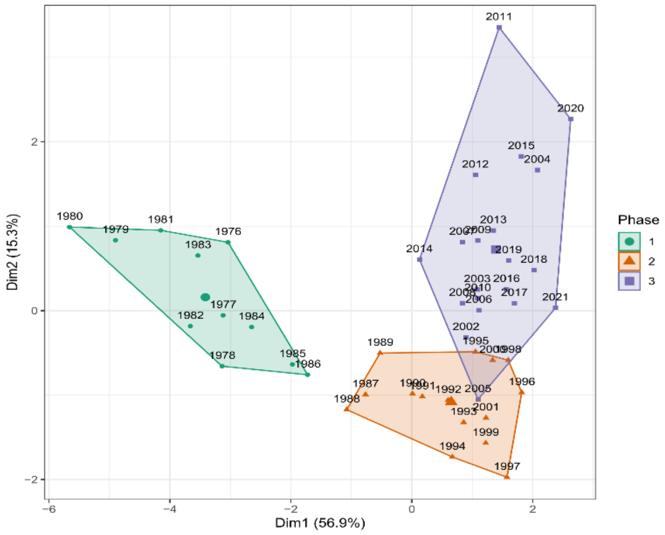


*Figure S 5: Cluster plot of fish community structure data based on the classification from GMM model-based clustering. Data points are coloured according to their assigned clusters, with convex hulls outlining each cluster.*

### 3. Data selection

To analyze the fish community structure drivers, we tested all environmental and zooplankton data in addition to fishing mortality of sprat, cod and herring. After variable selection using VIF/correlation check, we eliminated summer bottom temperature, which was highly correlated to winter bottom salinity. The resulting correlations between predictors are shown in Figure S6.

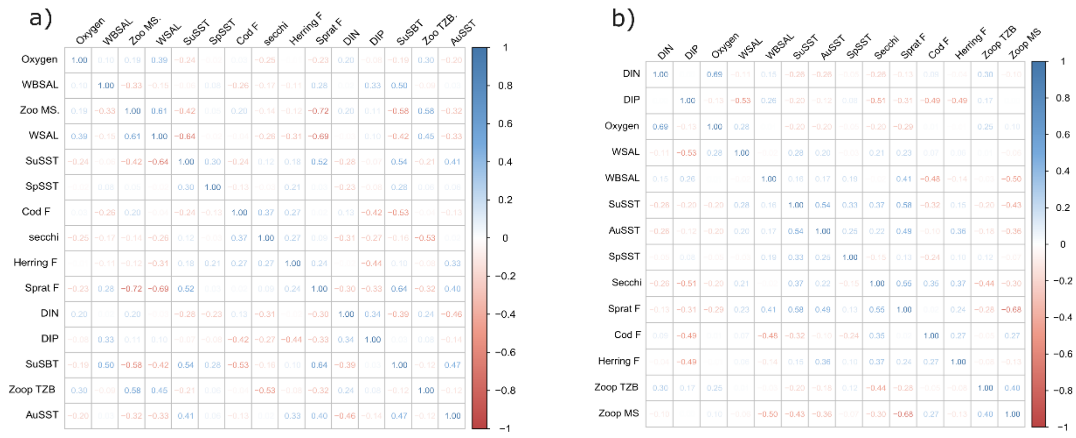


*Figure S 6: Correlation plot of predictors used for the fish community structure BRT analysis a) before and b) after the selection process. Prior to correlation analysis, Variance Inflation Factor (VIF) were applied for variable selections. The correlations chosen are <0.7.*

### 4. Boosted regression tree results

#### 3.1. Performance metrics

The BRT analysis was conducted for three distinct fish community structure phases,using the model settings: tree complexity = 5, learning rate = 0.001, and bag fraction = 0.75. The results include several performance metrics for both the training and cross-validation sets. The performance metrics include the Total Deviance (the total variation in the data), Residual Deviance (unexplained variation in the data after fitting the model), Correlation (strength of the relationship between the predicted and observed values), Area Under the Curve (AUC), Percentage Explained Deviance. The cross-validation (CV) statistics include CV Deviance, CV Correlation, CV AUC, and CV Percentage of Explained Deviance.

The BRT model performs well on training and cross-validation sets, with high correlation, AUC, and percentage of explained deviance. The model explains a large portion of the variance in the fish community structure in Supplementary Table 2 for all phases.

*Supplementary Table 2: Performance metrics and cross validation (CV) statistics for the fish community structure boosted regression tree (BRT) analysis. Tree complexity = 5, learning rate = 0.001, bag fraction = 0.75.*

| Performance metrics and cross validation | Phase 1 | Phase 2 | Phase 3 |
|---|---|---|---|
| Total Deviance | 1.10 | 1.26 | 1.37 |
| Residual Deviance | 0.02 | 0.04 | 0.02 |
| Correlation | 1 | 1 | 1 |
| AUC | 1 | 1 | 1 |
| Percentage of Explained Deviance | 98.14 | 96.92 | 98.72 |
| CV Deviance | 0.3 | 0.31 | 0.27 |
| CV Correlation | 0.88 | 0.95 | 0.93 |
| CV AUC | 0.97 | 1 | 1 |
| CV Percentage of Explained Deviance | 72.43 | 75.39 | 80.16 |

### 3.2. Partial dependency plots

Figure S7 shows the partial dependency plots that provide insights into the chosen predictors contribute to shifts in the system. These plots were used to identify the direction of the relationship for each predictor in different phases.

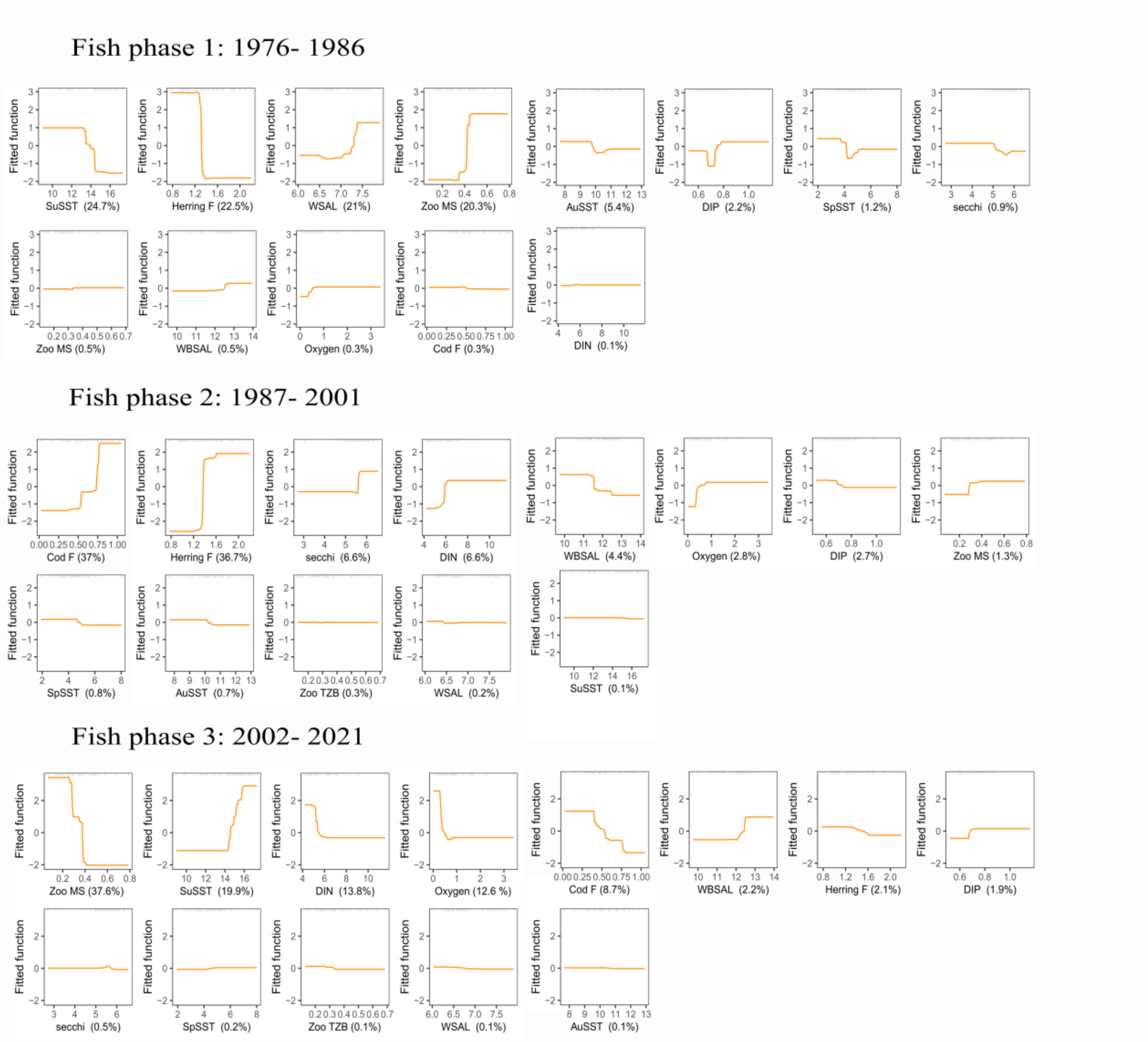


*Figure S 7: Partial dependency plots for predictors of each of the clusters for fish community structure. The graphs show the effect of a given predictor on the probability of occurrence of a shift while keeping all other variables at their mean. Relative influence of each predictor is reported between parentheses.*

#### 3.3. Fish BRT results without herring fishing mortality

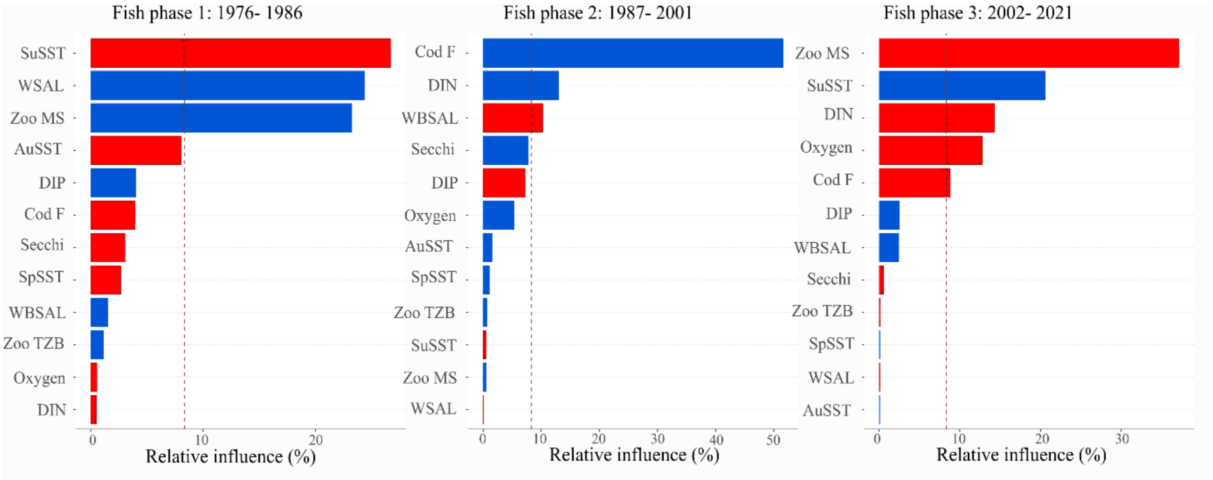


*Figure S 8: Analysis results of the fish community structure without herring fishing mortality. Relative influence of drivers/pressures during each of the phases identified with BRT analysis. The colors of the bars reflect the direction of the relationship with blue being positive, red being negative, and grey indicating a lack of clear direction. The dotted line indicate a relative influence above what could be expected by chance and considered as a driver (Jouffray et al. 2019).*

When taking away the herring fishing mortality, the driver identified as similar in the first and third phase but different in the second phase (Figure S8). In the second phase, we identify cod fishing mortality and dissolved inorganic nitrogen with positive directions of the relationship and winter bottom salinity (WBSAL) with a negative direction of the relationship. This indicates that lower bottom salinity contribute to driving the decrease in cod stock charactering the second phase. Please note that the performance metrics are lower than when counting herring fishing mortality as a driver although still acceptable (Supplementary Table 3).

*Supplementary Table 3: Performance metrics and cross validation (CV) statistics for the fish community structure boosted regression tree (BRT) analysis. Tree complexity = 5, learning rate = 0.001, bag fraction = 0.75.*

| Performance metrics and cross validation | Phase 1 | Phase 2 | Phase 3 |
|---|---|---|---|
| Total Deviance | 1.10 | 1.26 | 1.37 |
| Residual Deviance | 0.18 | 0.22 | 0.02 |
| Correlation | 0.97 | 0.97 | 1 |
| AUC | 1 | 1 | 1 |
| Percentage of Explained Deviance | 83.43 | 82.34 | 98.72 |
| CV Deviance | 0.7 | 0.7 | 0.39 |
| CV Correlation | 0.71 | 0.73 | 0.87 |
| CV AUC | 0.94 | 0.95 | 0.97 |
| CV Percentage of Explained Deviance | 36.27 | 44.38 | 71.41 |
